# An integrative model of leukocyte genomics and organ dysfunction in heart failure patients requiring mechanical circulatory support

**DOI:** 10.1101/024646

**Authors:** Nicholas Wisniewski, Galyna Bondar, Christoph Rau, Jay Chittoor, Eleanor Chang, Azadeh Esmaeili, Mario Deng

**Author notes:** Corresponding author: Nicholas Wisniewski,.

## Abstract

**Background:** The implantation of mechanical circulatory support (MCS) devices in heart failure patients is associated with a systemic inflammatory response, potentially leading to death from multiple organ dysfunction syndrome. Previous studies point to the involvement of many mechanisms, but an integrative hypothesis does not yet exist. Using time-dependent whole-genome mRNA expression in circulating leukocytes, we constructed a systems-model to improve mechanistic understanding and prediction of adverse outcomes.

**Methods:** We sampled peripheral blood mononuclear cells from 22 consecutive patients undergoing MCS surgery, at 5 timepoints: day -1 preoperative, and days 1, 3, 5, and 8 postoperative. Phenotyping was performed using 12 clinical parameters, 2 organ dysfunction scoring systems, and survival outcomes. We constructed a systems-representation using weighted gene co-expression network analysis, and annotated eigengenes using gene ontology, pathway, and transcription factor binding site enrichment analyses. Genes and eigengenes were mapped to the clinical phenotype using a linear mixed-effect model, with Cox models also fit at each timepoint to survival outcomes. Finally, we selected top genes associated with survival across all timepoints, and trained a penalized Cox model, based on day -1 data, to predict mortality risk thereafter.

**Results:** We inferred a 19-module network, in which most module eigengenes correlated with at least one aspect of the clinical phenotype. We observed a response to surgery orchestrated into stages: first, activation of the innate immune response, followed by anti-inflammation, and finally reparative processes such as mitosis, coagulation, and apoptosis. Eigengenes related to red blood cell production and extracellular matrix degradation became predictors of survival late in the timecourse, consistent with organ failure due to disseminated coagulopathy. Our final predictive model consisted of 10 genes: *IL2RA*, *HSPA7, AFAP1, SYNJ2, LOC653406, GAPDHP35, MGC12916, ZRSR2*, and two currently unidentified genes, warranting further investigation.

**Conclusion:** Our model provides an integrative representation of leukocyte biology during the systemic inflammatory response following MCS device implantation. It demonstrates consistency with previous hypotheses, identifying a number of known mechanisms. At the same time, it suggests novel hypotheses about time-specific targets.

## 1 Background

Mechanical circulatory support (MCS) device therapy is a treatment option for patients with advanced heart failure (AdHF), which consists of surgical implantation of a mechanical device to restore normal hemodynamics. It is commonly used as a bridge to heart transplantation for deteriorating patients awaiting an organ, as a bridge to recovery when myocardial function can still be restored, and as lifelong (destination) therapy for patients who are not candidates for transplantation [1]. The physiological rationale underlying the treatment is that by restoring normal hemodynamics, it is possible to at least partially restore systemic functions such as oxygen metabolism and neurohormonal regulation, which in turn reduces inflammation and improves the function of all organs and promotes the recovery of the heart failure patient [2]. However, surgical implantation of the device is associated with a dangerous systemic inflammatory response, which results in temporary acceleration of organ dysfunction, and potentially leads to death. It is presumed that an altered immune response is induced by both the MCS device and surgery, facilitated by the preexisting heart failure syndrome [3].

The most feared consequence of this systemic inflammatory response is multiple organ dysfunction (MOD) syndrome [4], the leading cause of mortality in intensive care units. Clinically, patients develop a cascade of complications, including hepatic, renal, pulmonary, immunologic, coagulation, gastro-intestinal, metabolic and neurological dysfunction. Few options exist to help reduce this risk, as most aspects of the progression to MOD are not well understood. Currently, the only clinical precaution is to place all patients on anticoagulants prior to surgery, in order to mitigate complications arising from abnormal coagulation. It is thought that activation of the immune-inflammation-associated coagulation pathway produces fibrin matrix that blocks blood flow and therefore causes tissue necrosis, ultimately leading to organ failure [5]. Interestingly, anti-inflammatory measures such as glucocorticoids have yielded mixed results [6], and are therefore not used.

A significant amount of work has been done examining similar therapies for systemic infection, i.e. sepsis–related MOD [7]. In both sepsis-and MCS-related MOD, the interplay between leukocytes, platelets, and endothelium is thought to play a critical role in mediating the phenotype [8, 9], and so many of the strategies are understood as affecting various aspects of the endothelial response. For example, a number of clinical trials have been conducted to evaluate antiadhesion therapies, anticoagulant therapies, antiapoptosis therapies; transcription factor targets such as NF-ĸB; signaling pathways such as MAPK; and nitric oxide synthase (NOS) inhibitors [10]. Unfortunately, most interventions have failed to improve outcomes, leaving much work to be done researching new therapeutic targets.

It is thus still believed that there are many interacting mechanisms in MCS-related MOD, *although there is currently no hypothesis about their integrative systems biology*. The coordination of all these features has yet to be contextualized, and many questions remain about how they are orchestrated together as a time-evolving system. Analysis of longitudinal whole-genome mRNA expression in circulating leukocytes offers a way to synchronize all of these known features, while also facilitating discovery of novel features. Several machine learning and bioinformatics tools are available to assist in statistical modeling of high-dimensional data, and have yielded promising results in many genomic studies. These methods can allow for discovery of interesting temporal features of the immune response and recovery process, and identify systemic motifs that are of clinical importance.

In this article, we profile whole-genome mRNA expression in circulating peripheral blood mononuclear cells (PBMC). PBMC is commonly used to study leukocyte gene expression because it is easily accessible and includes several key inflammatory cell populations. To reconstruct the temporal dynamics in response to surgery, we analyzed a short time-series of samples taken at 5 time points during the first week, including day -1 preceding surgery. Because most of the dynamic response to surgery occurs in the first postoperative week, it is expected that patient trajectories are most sensitive and informative during this window. Our analysis strategy then uses machine learning to gain a comprehensive picture of the systems biology and its relation to clinical outcomes.

## 2 Methods

### 2.1 Patients

We collected blood samples from 22 consecutive AdHF patients undergoing MCS between March 2010 and September 2014 at a single institution. Samples were collected at 5 timepoints: day -1 preoperative, and days 1, 3, 5, and 8 postoperative.

### 2.2 Clinical management

All study participants were referred to the UCLA Integrated Advanced Heart Failure Program and evaluated for the various therapeutic options, including continued optimal medical management, MCS, and heart transplantation. All study participants were recommended by the multidisciplinary heart transplant selection committee to undergo MCS-surgery therapy, and consented to proceed. Preoperatively, n=2 patients were in a state of critical cardiogenic shock (Interagency Registry for Mechanically Assisted Circulatory Support (INTERMACS) class 1); n=12 patients were progressively declining despite being on inotropic support (INTERMACS class 2); n=7 patients were stable but inotrope dependent (INTERMACS class 3); and n=1 patient had resting failure symptoms (INTERMACS class 4) [1]. All patients were optimized regarding medical heart failure therapy, and were undergoing MCS-therapy according to established guidelines [11, 12].

After anesthesia induction, patients were intubated and placed on cardiopulmonary bypass. The type of MCS-device selected depended on the acuity and severity of the heart failure syndrome as well as patient characteristics [13]. For left ventricular support, patients underwent either Heartmate II (HeartMate II® pumps are valveless, rotary, continuous flow pumps) or HVAD (HeartWare® HVAD pumps are valveless, centrifugal continuous flow pumps). For biventricular support, patients underwent either Centrimag-BVAD (Centrimag® pumps are valveless, centrifugal, continuous flow pumps that are external to the body), PVAD-BVAD (Thoratec® Paracorporeal Ventricular Assist Device (PVAD) pumps each contain two mechanical tilting disk valves) or the t-TAH (the Temporary Total Artificial Heart consists of two artificial ventricles that are used to replace the failing heart).

Various combinations of cardiovascular stimulant drugs were used to support patients postoperatively, tailored to individualized requirements. In addition, other temporary organ system support was administered as required (e.g. respirator, dialysis, transfusions, antibiotics).

### 2.3 Clinical phenotyping

We collected 12 distinct parameters on a daily basis for detailed clinical phenotyping of the patient cohort. At each timepoint, the following clinical parameters (if present) were recovered from patient records:

1. Bilirubin
2. Creatinine
3. Leukocyte count
4. Platelet count
5. PaO2/FiO2 ratio
6. Mean arterial pressure
7. INR (International Normalized Ratio, for prothrombin time)
8. Glucose
9. Heart rate
10. Respiratory rate
11. Temperature
12. Glasgow Coma Scale (GCS)

Using combinations of these parameters, we also computed two validated and commonly used composite organ dysfunction scores, SOFA [14] and MELD-XI [15]. The SOFA score is a validated and widely accepted measure that rates degree of organ failure across 6 major organ systems (cardiovascular, respiratory, neurological, renal, hepatic, and coagulation). The MELD-XI score uses only the bilirubin and creatinine levels.

Additional information can be found in the Appendix.

### 2.4 Sample processing

#### 2.4.1 Sample collection and RNA isolation

Mononuclear cells were isolated from 8 ml of blood collected by Vacutainer cell preparation (CPT) tubes with sodium citrate (Becton Dickinson, Franklin Lakes, NJ), resuspended in RNA Protect (Qiagen, Valencia, CA) within 2h of phlebotomy. Total RNA was isolated from each sample (RNeasy, Qiagen, Valencia, CA). Quality of the purified RNA was verified on an Agilent® 2100 Bioanalyzer (Agilent Technologies, Palo Alto, CA); RNA concentrations were determined using a NanoDrop® ND-1000 spectrophotometer (NanoDrop Technologies, Wilmington, DE).

#### 2.4.2 RNA processing and analysis

After RNA extraction, quantification and quality assessment, total mRNA was amplified and hybridized on the Illumina HiSeq2000 TruSeq. Data was then subjected to quantile normalization using GenomeStudio (Illumina, San Diego, CA). Batch effects were removed using the ‘ComBat’ algorithm in R [16].

Prior to analysis, the entire set of mRNA transcripts (39,740) was filtered by variance and entropy criteria to remove uninformative transcripts. Filtering has been demonstrated to improve the interpretability of the inferred co-expression network, as well as reduce the bias in cluster analysis [17]. We therefore first removed genes with zero variance, and then placed cuts on the remaining transcripts to remove the remaining lower 30^th^ quantiles, reducing our list of transcripts to 14,753.

### 2.5 Statistical analysis

We began our analysis by using unsupervised machine learning to generate a systems representation. We used weighted gene co-expression network analysis (WGNCA) [18] to cluster co-regulated gene modules, construct an eigengene network, and compute network properties for each gene. The eigengene approach offers a powerful way to reduce dimensionality, while maintaining many useful statistical properties and biological interpretability. The result is a minimally biased systems-representation, providing a useful framework for relational and mechanistic reasoning.

After the network was constructed, we inferred each module’s clinical relevance using a linear mixed-effect model, relating the module eigengenes to the panel of day-to-day clinical parameters. The linear mixed-effect model accounts for the repeated-measures structure, and at this stage we controlled for confounding variables (e.g. age, sex, race, device, diabetes, ischemic etiology, platelet transfusions, plasmapheresis, and immunosuppression).

We inferred the biological relevance of each module using common bioinformatics tools: gene ontology (GOSim [19]), pathway (Strand NGS [20]), and transcription factor binding site (rVista [21]) enrichment analyses. These tools allowed us to investigate each module for over-represented groups of genes with known associations, providing the connections to biological processes necessary for interpreting the eigengenes.

To infer temporal patterns, we scaled the median eigengene levels along their temporal aspect, and sorted each temporal slice by z-score to identify salient features. To infer which eigengenes were related to survival outcomes, we used a multivariate Cox mixed-effect model to look for overall module effects, as well as separate univariate Cox models at each timepoint.

When working with unsupervised methods such as WGCNA, there is always concern that the eigengenes may be too great an abstraction, and that important statistical properties have been compressed away. To verify that the eigengene model is consistent with the statistical properties of the genes themselves, we computed models for each individual gene, analogous to our analysis of eigengenes: we used a linear mixed-effect model to infer relationships between genes and clinical parameters, and we used a Cox model at each timepoint to infer associations between genes and survival outcomes. We then tested the hypothesis that predictive eigengenes are associated with predictive genes. A hypothesis of no association implies that the distribution of p-values within a module should be flat, while a true association will exhibit an abundance of statistically significant genes that skew the p-distribution. To quantify this hypothesis test, we analyzed the distribution of p-values for genes in each module using a binomial test, performed by dichotomizing around the usual significance threshold *p*=0.05.

Finally, there is general interest in predictive models involving a very small panel of genes that can be easily and quickly sampled in typical clinical applications. These models care little for interpretability, aiming only for accuracy. The simplest model to consider would be stationary, with the same parameters at any timepoint – a single predictive tool that can be applied at any time. For this purpose, we created a low-dimensional predictive model by selecting only genes that achieved a Cox significance of *p*<0.1 at all five timepoints. Then, because of multicolinearity, we trained a multivariate Cox model using an elastic-net penalty (mixing parameter α=0.1). In the absence of a separate testing cohort, we adopted the following strategy: we trained the model on the day -1 pre-surgery samples, optimized the penalty to minimize error under 10-fold cross-validation, and tested on post-surgery samples. This approach is still biased in that we used univariate p-values across all timepoints as a filter to choose the predictors that went into the day-1 model. But the testing is able to assess the stability of the multivariate coefficients, since they were trained on only the day -1 timepoint; success in predicting survival outcomes from the subsequent postoperative data indicates stability of the coefficients over time.

Additional information can be found in the Appendix.

## 3 Results

### 3.1 Clinical phenotyping

Main demographics of the study patient cohort are summarized in Table 1. Of 22 patients undergoing MCS-surgery, 5 patients died on the MCS device at 32 (20-50) days after surgery, where we report results in the usual format of median (interquartile range). The 17 survivors were followed until transplant (n=13) for 107 (61-220) days or follow-up end (December 31, 2014) (n=4) for 600 (260-799) days. All non-survivors were male, and a significantly higher fraction of non-survivors had an underlying ischemic etiology.

We studied the main temporal clinical characteristics from Day -1 to Day +8 (Fig. 1). As a first visualization, we used principal component analysis, and charted a median timecourse (Fig. 1A). Similar parameters have the same orientation, e.g. MELD-XI, SOFA, creatinine and bilirubin cluster together. Anti-correlated variables have opposing orientations. Note that there is a clear separation between medians of survivors and non-survivors along the SOFA and MELD-XI dimensions. This is expected, as non-survivors have overall worse organ dysfunction scores.

We next examined the median timecourse of each individual parameter (Fig. 1B). On the day following surgery, we see the SOFA score peak, while the platelet count, temperature, and respiratory rate trough. The white blood cell count reaches its maximum on day +3, and then recovery occurs on day +5 and +8 as the platelet count rises and the SOFA score decreases.

Finally, we fit univariate Cox models on each clinical parameter at each timepoint to identify parameters predictive of survival (Fig. 1C). We found that the organ dysfunction scores (SOFA and MELD-XI), driven by bilirubin and creatinine, became more predictive of survival at later timepoints. This is expected, as the survivors improve over time, but the non-survivors do not. We also see platelet count as a predictor of outcome both early and late in the timecourse. This observation supports the clinical practice of administering platelet transfusions to patients with low platelet counts. Also, we noticed that the survivors tended to increase platelet count during the recovery stage, while non-survivors didn’t.

### 3.2 Eigengene analysis

#### 3.2.1 Eigengene network

We inferred a network that consisted of 19 gene modules. One of the WGCNA modules is always designated as a leftover module for genes that went unclustered, and we labeled it as such. We then labeled the remaining modules with biologically relevant terms using gene ontology enrichment analysis (Fig. 2A), which we report in full in Table Supplement 3. We next did pathway analyses for each module, and enrichment analysis of transcription factor binding sites, both of which are summarized in Table Supplement 4. An in-depth discussion of the systems biology is in the Discussion section.

To reduce the complexity even further, we looked for biologically interpretable superclusters by hierarchically clustering the module eigengenes. We identified 5 superclusters, and color-coded them (Fig. 2A). Much of this structure can be explained succinctly: the turquoise supercluster can be understood as the adaptive immune system, the brown supercluster as components of the innate immune system, blue as metabolic, green as catabolic, and yellow as a reparative supercluster. These supercluster colors are used throughout to aid in interpretation and organization.

To gain a more integrated understanding of the network, we computed correlations between eigengenes and analyzed the eigengene network (Fig. 2B). The most distinctive feature of the network is the strong negative correlation between the innate and adaptive immune superclusters. This feature has been previously commented on in the literature [22]. We also find strong correlations stemming from the mitochondria module, situated between these opposing superclusters. Its position suggests a close working relationship between the mitochondria and innate immunity modules.

This hypothesis is also supported by the literature. Mitochondria have been considered principal mediators of inflammation and arbitrators of the pro-inflammatory state [23], have been characteristically studied in the context of organ dysfunction, and play a critical role in initiating tissue hypoxia and production of reactive oxygen species [24-26]. Mitochondrial products have also been proposed as damage-associated molecular pattern (DAMP) signals [27] [28]. As we will see in the time-dependent analysis, the mitochondria and innate immune eigengenes peak together within the first 24h after surgery. This coincidence suggests temporally orchestrated interactions between the modules.

#### 3.2.2 Mapping eigengenes to clinical parameters

The next step was to connect the eigengenes to the clinical phenotype. We inferred the clinical relevance of each module using a linear mixed-effect model. We then identified patterns by bi-clustering the eigengenes with the clinical parameters using the –log p-values from the model (Fig. 3).

Organ dysfunction, as measured by the SOFA and MELD-XI scores (along with creatinine, glucose, and white blood cell counts), was positively associated with the innate immunity and metabolic superclusters, particularly the innate, metabolic, and mitochondria modules. These organ dysfunction parameters were negatively associated with the adaptive immunity supercluster, particularly the T cell, B cell, NK cell, Type I IFN, demethylation and transcription modules. This is consistent with previously reported innate immunity activation and T-cell suppression [22]. As innate-immunity-associated inflammation increases, organ dysfunction worsens, and the innate immune system suppresses the adaptive immune system. As inflammation is suppressed, organ dysfunction improves, and the adaptive immune system function returns.

The clustering of glucose levels with the organ dysfunction scores is also consistent with the literature. Higher glucose levels have been associated, likely via increased adrenergic and cortisolergic drive, with an increased 30-day mortality rate in conditions such as acute heart failure [29, 30].

We also noted these superclusters had significant associations with the platelet count, which is a parameter that contributes to the SOFA score, but through an inverse relationship. We therefore see the opposite of the SOFA associations: the platelet count had a positive association with the T cell, NK cell, type I IFN, and coagulation modules, and a negative association with the metabolic, innate, and mitochondria modules.

#### 3.2.3 Time-dependent eigengene analysis

We expected to see distinctive time-dependent features in the transcriptome, because the surgery is a significant perturbation to the system, and the recovery is very dynamic. As with the clinical parameters, we characterized these features by examining how the median eigengene levels changed over time.

As a first visualization, we projected the timepoint medians onto the leading principal components of the eigengene representation, and charted a median timecourse (Fig. 4A). The orientation of the eigengenes reflects the superclustering we have previously noted, where we note the opposition of the innate and adaptive immune system, with the remaining eigengenes acting orthogonally to both. We notice a net displacement from day -1 to day 8 in the direction of the reparative and catabolic superclusters, indicative of the recovery processes activated following the surgery. The second timepoint, however, is a large deviation from that trajectory, which we associate with activation of the innate immunity supercluster, and suppression of the adaptive immunity supercluster.

Next, we examined the median timecourse of each eigengene separately to better identify time-dependent features. We converted the median eigengene levels to standard z-scores, where the scaling was done along the time aspect, and ranked within temporal slices to identify salient features at each time point. This process is illustrated in Figure 4B, and we summarized the temporal eigengene features in Table 2. A more in-depth explanation of the result is included in the Discussion.

Finally, to infer which eigengenes are related to survival outcomes, we used a multivariate Cox mixed-effect model to look for time-averaged effects. Anticipating dynamic effects, we also fit separate univariate Cox models at each timepoint. The results of both are shown in Figure 4C. We notice a consistent effect from the demethylation module across all timepoints. The demethylation module is heavily enriched by Y-chromosome genes, indicating that this effect is likely due to the fact that all non-survivors were male. We also notice mitosis and defense eigengenes becoming predictive late in the timecourse. These later features are related to red blood cell production and extracellular matrix (ECM) degradation, and are consistent with the hypothesis that organ failure is caused by fibrin matrix blocking blood flow leading to necrosis.

### 3.3 Eigengene statistical properties

To verify that the eigengene model is consistent with the statistical properties of the genes themselves, we fitted separate models for each gene, directly analogous to our analysis of the eigengenes. We then tested the hypothesis that predictive eigengenes are associated with predictive genes, by analyzing the distribution of p-values in each module using a binomial test performed by dichotomizing around p=0.05. The distribution of predictive genes is consistent with the predictive patterns found throughout the analysis of the eigengenes, including the time-dependent patterns (Fig. 5). This result demonstrates that the eigengene-phenotype associations we inferred are representative of the individual gene-phenotype associations.

### 3.4 Clinical risk prediction

Finally, we created a low-dimensional predictive model by selecting only genes that achieved a univariate Cox significance of *p*<0.1 at all five timepoints. Only eleven genes passed this requirement. Because of multicolinearity, we performed the Cox regression with an elastic-net penalty (mixing parameter alpha=0.1), which had the effect of driving one of the coefficients to zero while adjusting the rest, leaving a ten-gene model (Fig. 6A). In the absence of a suitable testing cohort due to our small sample size, we trained the model on the day -1 preoperative samples, optimized the penalty to minimize error under 10-fold cross-validation, and tested on post-surgery samples. In this way, the coefficients were only determined using the preoperative data, and success in predicting survival outcomes from the postoperative data indicates stability in the coefficients over time. We found our coefficients to be stable: they demonstrated the ability to distinguish survivors from non-survivors, when tested on the postoperative data, with perfect accuracy. This suggests that a single model, which works at all timepoints, is feasible for predicting MOD outcomes.

We examined the module memberships of each gene in the model, and looked for patterns by clustering (Fig. 6B). We found that half the genes, *SYNJ2, IL2RA, AFAP1, MGC12916* and *HSPA7*, have strong associations with the adaptive and innate superclusters. We examined the association of each gene with the clinical parameters (Fig. 6C), and bi-clustered the signed –log p-values (the sign comes from the linear mixed-effect model coefficient), to look for patterns. We found that *NEWGENE532* has very strong associations with the organ failure scores, and *ZRSR2* has a very strong association with PaO2/FiO2.

The annotated genes in the ten-gene predictive model are involved in a variety of processes. *Afap1,*a potential modulator of actin filament integrity in response to cellular signals, and *Synj2* which has been marked as a candidate gene for overall life span [31] are known mediators of cell division while the splicing factor *ZRSR2* plays a critical role in the development of blood cells (RBC, WBC and platelets) from the bone marrow [32]. Since an important distinguisher for survival is the replenishment of lost blood cells (especially platelets) on days 5 and 8, the presence of these genes is expected.

Equally important to the replenishment of lost cells is the maintenance of living cells. *HSPA7* is a member of the *HSP70* chaperone protein family, and helps mediate cellular apoptosis under stress conditions, and is of importance in pathological conditions in that it contributes to cell survival [33]. *Il2ra* is a regulator of regulatory T-cell function. Regulatory T-cells act to modulate the response of the immune system to distress, and play an important role in the prevention of autoimmunity. Taken together, the annotated genes suggest that maintenance and replenishment of cells are crucial to the survival during MOD.

The roles of the other genes in the predictive model are largely unknown, with three (*LOC653406, GapdhP35, Mgc12916)* with poor annotations and two (*NEWGENE532 and NEWGENE15*) having no annotation whatsoever. *NEWGENE15* is located in the T Cell module, and likely plays a role similar to the other 4 genes in the predictive model found within that module. Regardless, further analysis of all of the genes in the predictive model is warranted, and we will include the mRNA sequences of the new genes in the final publication.

## 4 Discussion

We now present a detailed discussion of our investigation into each module’s role in the systems biology of the systemic inflammatory response, organized by superclusters.

### 4.1 Innate immune supercluster

#### 4.1.1 Innate

The innate immune system initially responds to the surgical intervention by the activation of a variety of immune cells including basophils, neutrophils, macrophages and eosinophils. It generally provides a preliminary response to a pathogen by activation of toll-like receptors (TLRs) via pathogen-associated molecular pattern molecules (PAMPs), or damage-associated molecular pattern molecules (DAMPs), which are cell-derived and initiate and perpetuate immunity in response to trauma, ischemia, and tissue damage [34]. In our pathway analysis of this module, hubs involved in the enriched pathways include inflammatory cytokines such as IL-1 and TNF-α. The activation of these cytokines results in activation of signal transduction pathways (like RAC1), stimulating further production of cytokines, procoagulants, reactive oxygen species, and proliferative factors.

As expected, the innate module spikes on day 1 after surgery, initiating the inflammatory response (Fig. 4B). This is associated with worsening organ dysfunction, as measured by the SOFA and MELD-XI scores (Fig. 3). The innate module is also associated with higher blood glucose levels, which is not surprising given the overlap with the mitochondria module. It is also associated with higher white blood cell count (Fig. 3), which has a delayed peak on day 3 (Fig. 1B). Although the Cox model was not statistically significant at any timepoint, the trend of the coefficients suggests a more extreme activation on day 1, and a more extreme supression on day 3, in non-survivors compared to survivors (Fig. 4C).

#### 4.1.2 Apoptosis

The apoptosis module is correlated with the metabolic module, and anti-correlated with the transcription module (Fig. 2). Like the innate, metabolic, and mitochondria modules, it is associated with higher blood glucose (Fig. 3). It is enriched for signaling and cytokine activity, indicating that mitochondrial apoptotic pathways are being activated by pro-inflammatory cytokines. These pathways are known to play in important role in organ dysfunction, because the termination of the inflammatory response is effected through apoptosis [35]. In patients with sepsis, neutrophil apoptosis is inversely proportional to the severity of organ dysfunction [36]. We further observe that the module is associated with granulocyte specific cell markers, and shows a strong (*p*=8.4E-9) enrichment for genes associated with decreased susceptibility to endotoxin shock, suggesting that this module may be regulating the above pathway. We found it inhibited through most of the timecourse, and upregulated primarily on day 8 when the innate response and corresponding organ dysfunction are at a minimum (Fig. 4B), although there are hints of association to survival on day 3 when we see the innate response begin to terminate (Fig. 4C).

#### 4.1.3 Metabolic

The metabolic module’s phenotypic (Fig. 3) and temporal profile (Fig. 4B) is indistinguishable from the innate module, and they share similar correlations in the eigengene network (Fig. 2B). In contrast to the innate module’s Cox profile, the metabolic module shows a consistent effect on survival outcomes across postoperative timepoints, with levels in non-survivors being elevated (Fig. 4C). One difference in the make-up of these two modules is the type of immune cell markers that are observed within each module. An examination of the 100 genes with highest intramodular connectivity within each module reveals that the innate immunity module is enriched (*p*=.0016) for monocyte markers while the metabolic module is enriched (*p*=.011) for neutrophil markers. Neutrophils have previously been related to organ dysfunction [8], and may explain the difference in Cox profile compared to the monocyte-rich innate immunity module.

### 4.2 Metabolic supercluster

#### 4.2.1 Demethylation

The demethylation module is seemingly unconnected to the rest of the eigengene network (Fig. 2B). The module is filled with Y-chromosomal genes (Benjamini-Hochberg corrected *p*=3.3E-11) and X chromosomal genes (Benjamini-Hochberg corrected *p*=3.9E-4). Upon inspection of intramodular connectivity, we find that genes involved in X-inactivation (*Xist* and *Tsix*) are hubs of this module. However, we believe this module to be an artifact of our study design. WGCNA seeks to find groups of genes whose expression varies in a similar way across samples. Y-chromosomal and X-inactivation genes are only expressed in men and women, respectively. Therefore, clear expression patterns for these genes will be detected by WGCNA, and these genes will be grouped together into a single module. We find enrichment for H3K4 demethylases, likely due to the fact that there are very few of these demethylases, and one each is located on the Y and X chromosomes. In the Cox model, we find it to be consistently predictive of survival across all timepoints (Fig 4C), but given that all non-survivors in our study were male, we find it likely that this is the proper explanation for the association.

#### 4.2.2 Coagulation

Clinical coagulation management to prevent bleeding (i.e. platelet transfusions) and clotting (i.e. anticoagulation medication), is considered important to the survival of patients with MOD [9]. The coagulation module shows a strong correlation with platelet count, both overall (Fig. 3) and temporally (Figs. 1B and 4B). This module shows strong enrichments for coagulation and platelet aggregation, and shows cell-type specificity for megakaryocytes. We further observe that several known repressors of clot degradation are present in this module as well. When the top 100 genes most significantly associated with the SOFA score are examined, we also observe enrichment for mitosis and cell division pathways, suggesting that the module’s effects are likely due to platelet maturation.

#### 4.2.3 Mitochondria

The mitochondria module is strongly associated with worsening organ dysfunction (Fig 3), and it is temporally synchronized with the innate immune response (Fig. 4B). This has previously been noted in the literature. Several platelet mitochondrial respiratory-chain enzymes are known to be inhibited during human sepsis and organ dysfunction, e.g. cardiogenic shock [37], and mitochondrial biogenesis is associated with survival in MOD [25, 38]. Pathway analysis revealed enrichment of the PAR/Thrombin pathway, which helps initiate coagulation. The activation of this pathway signals production of ROS in the mitochondria, and its activation after MCS surgery may contribute to adverse outcomes [39]. Pathway analysis revealed relationships to hub genes such as cytokines IFN-γ and IL-8, as well as the signal transduction gene MAPK1. These may indicate processes associated with macrophage activation, as well as recruitment of other innate immune cells (IFN-γ and MAPK1 activate macrophages, while IL-8 stimulates trafficking of other cells to site of infection). Another pathway hub identified with the mitochondria module is BLC2, which has a role regulating apoptosis [26].

#### 4.2.4 Ribosome

The ribosome module upregulates immediately following surgery, and remains activated throughout recovery (Fig. 4B). As the body recovers from surgery, robust protein translation is an important component of many vital cellular processes, including cell division and the response to external stimuli and stressors. Furthermore, ribosomal proteins may play a protective role in host immune response by boosting immune signaling [40]. In the Cox profile (Fig. 4C), we observe underexpression in non-survivors on day 1, followed by overexpression on day 3.

### 4.3 Catabolic supercluster

#### 4.3.1 Protein folding

The protein-folding module shares a similar time profile with the ribosome module (Fig. 4B). It is strongly activated on day 8 along with the coagulation, catabolism, and apoptosis modules, but is also mildly upregulated on the day after surgery. Unlike the ER module, which appears to be enriched for stress-related chaperone proteins, this module is enriched for *de novo* posttranslational protein folding-associated genes. Additionally, there is a slight enrichment for genes involved in cytoskeleton maintenance and organization, suggesting that this module may be involved in the creation and maintenance of new cells after the catabolic processes have finished.

#### 4.3.2 Catabolism

The catabolism eigengene is negatively associated with the SOFA score (Fig. 3) and in many ways behaves similar to the coagulation module in terms of its relevance to MOD. Like the coagulation module, it is progressively activated throughout the study timecourse. It also appears to decrease mortality risk on day -1 and day 5 (Fig. 4C). Because a relevant therapy for the prevention of MOD mortality is the use of anti-clotting agents such as warfarin after surgery has ended, our initial assumption was that this module would correspond to this function. To our surprise, none of the major members of the clot removal cascade are found in the module; there are very few ECM remodeling genes present. Instead, examination revealed a number of macrophage, autophagy and apoptosis-related genes, suggesting that the module is involved in the ubiquitination cascade and whole-cell turnover. We examined the top 100 genes in the module based on intramodular connectivity, and observe that these important genes are enriched for apoptosis and the adaptive immune response, as well as enrichment for genes expressed in proerythrocytes and genes that are causal for premature red cell death and anemia. Put together, these data suggest that this module is involved in the maintenance of blood homeostasis, ensuring that a proper amount of erythrocytes are produced in the body and acting to eliminate excess RBCs and WBCs that are produced. This matches well with the final predictive model, which included several genes involved in blood homeostasis and cell division.

#### 4.3.3 RNA processing

The RNA processing eigengene is correlated with the catabolism eigengene (Fig. 2B). RNA processing is an important component of the cell growth, division and response to external stressors. Its activation on the final two days suggests that as the body recovers from surgery, the reparative processes are being robustly activated (Fig. 4B).

### 4.4 Reparative supercluster

#### 4.4.1 ER (Endoplasmic Reticulum)

The ER eigengene rises continually after surgery (Fig. 4B) and shows a strong enrichment for the ER unfolded protein response, suggesting that this module is involved in the management of the intracellular reparative response after surgery.

#### 4.4.2 Mitosis

The mitosis module is strongly enriched for genes involved in cell cycle maintenance and mitosis in general, and shows enrichment for markers of prorythrocytes, the precursor cell of erythrocytes. This is notable because of the enrichment seen in the catabolism module for erythrocyte cell markers, and the genes in the predictive model that correspond to blood homeostasis. The mitosis module is correlated with the catabolism module (Fig. 2B), and peaks at day 5 (Fig. 4B). The Cox model identifies it as associated with survival on days 5 and 8 (Fig. 4C). The late-stage activation of this module suggests that mitosis of erythrocytes, as well as platelets (as in the coagulation module), are important to repair and survival with MOD.

#### 4.4.3 Defense

The defense module shares several features with the mitosis, coagulation and catabolism modules. Like these modules, it is progressively activated over the course of the study (Fig. 4B), and like the mitosis module, it is predictive of survival late in the timecourse (Fig. 4C). It shows a significant association with worsening bilirubin (Fig. 3), and enriched for genes involved in the innate immune system, specifically granulocytes and neutrophils. Neutrophil dysregulation has previously been associated with negative outcomes in MOD [8]. More broadly, granulocytes act to regulate inflammation in the body. Additionally, this module shows enrichment for a number of ECM remodeling proteins, suggesting that the granulocytes are assisting in the degradation of platelet clots (whose degradation is suggested by the beneficial role of anticoagulants in treatment of MOD), the freeing of dying cells from the ECM, or negatively affecting the integrity of the endothelial cell wall.

### 4.5 Adaptive immune supercluster

#### 4.5.1 B cells

The activity of the B cell module follows an interesting timecourse: it peaks on day 3 after surgery, in an intermediate phase between the innate immune response and the adaptive immune response (Fig. 4B). The Cox model shows a very slight, but consistent, effect on survival across all timepoints (Fig. 4C). One possible explanation is that non-survivors experience upregulation of plasma B cell subsets which are characteristically known to create further inflammation. We also find greater upregulation of IL-12B receptors in the non-surviving patients, across all timepoints. IL-12 is a critical regulator of the cell mediated Th1 response, and therefore is an important promoter of autoimmunity and inflammation. Finally, non-survivors may fail to express enough anti-inflammatory IL-10 immediately after surgery. IL-10, which is secreted by regulatory B cells (Bregs), plays a tremendous role in regulating the adaptive immune system and the inflammatory responses [41]. Immediately following surgery, we found that Il-10 is more highly expressed in survivors, suggesting that immediate Breg activity may be essential to survival post-surgery. Depending on the complex interplay between the B cell subsets, the inflammation response can hasten or subside, thereby drastically affecting recovery after MCS surgery.

Many genes in this module are associated with B cell activation and proliferation, such as CD19, MS4A1 (CD20), and CD22. CD20 and CD22 have been studied in context of inflammation and autoimmune diseases [42]. Furthermore, pathway analysis revealed enrichment in signal transduction through the B cell and Par 1 receptors. The high expression of genes in this module may indicate that the activation of B cells in particular are propagating the inflammation, and even assist in instigating coagulation processes via thrombin/Par1 pathway. However, another important factor in interpreting this module is the potential presence of Bregs, which mediate several anti-inflammatory events [41].

#### 4.5.2 Transcription

Like the B cell module, the transcription module peaks on day 3 (Fig. 4B). This module is anti-correlated with the apoptosis, metabolic, innate, and mitochondria modules (Fig 2B). Since the innate, apoptosis and metabolic modules are associated with increased innate immune cell function, and the mitochondrial module is strongly associated with organ dysfunction (Fig 3), it seems likely that the transcription that is occurring in this module is linked to the role of the adaptive immune system in regulating the response of the body after the initial inflammatory phase is completed. Robust gene expression changes are important for proper responses to stressors and external stimuli, and the anti-correlation of this module to the four modules listed above suggests that the genes being transcribed are acting beneficially.

#### 4.5.3 T cells

The T cell module is the largest module in the adaptive immune supercluster (Fig. 2A), and is the module most strongly associated with improvement of organ dysfunction (Fig. 3). Its expression is highly suppressed during the initial innate immune response, and then it steadily increases over the timecourse (Fig. 2B), exhibiting a statistically significant association with the platelet count (Fig. 3). The Cox model shows a mild negative association with survival, along with the NK cells and B cells, contrary to the association with improving organ failure scores (Fig. 2C). Although this association with survival did not achieve statistical significance, it is fairly consistent across timepoints, suggest the effect is real. This is not necessarily paradoxical, as high levels of T cells are the result of the body’s attempt to deal with advanced disease. There may be a simple analogy to explain this result: firefighters put out fires, but the presence of a large number of firefighters at a given fire is indicative of a fire too large to control. T cells have previously been reported to play an important role in the regulation of organ failure, with the depletion of peripheral blood CD4+ T-cells being associated with persistent organ failure [43]. T cells have also been consistently shown to play an important role in inflammation-based organ damage [44]. Furthermore, these cells play a vital role in transplant rejection by mediating T-cell induced rejection [45]. Taken together, the proper regulation of this module acts to suppress organ dysfunction.

#### 4.5.4 NK cells

The NK cell module has a similar profile to the T cell module. Like T cells, it is highly suppressed during the innate immune response, and continually rises throughout the rest of the timecourse (Fig. 4B). It also shares a similar Cox profile with the T cell module (Fig. 4C). NK cells play an important role in assisting T cells in host defense [46], and when improperly regulated, may result in widespread organ failure [47] and auto-immune disorders [48]. It likely acts in concert with the T-cell module to modulate the immune response to the surgery and, when properly controlled, protects the host from multi-organ dysfunction.

#### 4.5.5 Type I IFN (Interferon)

The type I IFN module contains pathways and genes related to signal transduction—particularly with interferon, leukocyte proliferation, and leukocyte trafficking in both innate and adaptive processes. It shares pathways with the innate immune system, but is temporally anti-correlated with the innate response (Fig. 4B). The module is heavily downregulated after surgery, and then increases expression at the later timepoints. This module becomes predictive of survival on the final day 8 (Fig. 4C), confirming the previously described potential of type-I IFN to restore immunocompetence and improve survival outcomes [49]. Considering this is a later stage in the time course, and that genes in this module tend to be enriched (*p*=.011) for helper T-cell markers, this module may be associated with T cell function.

### 4.6 Study design discussion

Providing the best possible care for advanced heart failure patients requiring advanced therapies is an important challenge for modern medicine. The mortality rate following MCS implantation is relatively high compared to other medical procedures, in the range of 20% [1]. Understanding how to better understand and predict clinical outcomes requires a model that accurately captures a multitude of complex features in the systemic inflammatory response.

The necessity of developing an appropriate conceptual immunological framework has been stated by various groups [24, 50], but has been accompanied by significant challenges. Genome-wide transcription profiling in human sepsis has indicated that both pro-and anti-inflammatory mechanisms occur at various times over the course of sepsis [34, 51], and that a patient may cycle through each phase multiple times [52]. This indicates fundamental problems for the classical biphasic model, which assumes a systemic inflammatory response syndrome (SIRS) followed by a compensatory anti-inflammatory response syndrome (CARS) [53]. However, recent investigations continue to build upon this limited model, stating that the biphasic view “may be a simplistic explanation of a complex disease, yet provides a rational explanation for how the function of the immune system becomes altered during the course of sepsis” [54]. An integrative model that moves beyond the current SIRS/CARS framework would be of great use to the study of MCS-and sepsis-related MOD.

In this study, we developed an integrative model by utilizing a systems biological approach, based on three research study design assumptions. First, we take the clinical phenotypic trajectory as the most reliable, relevant, and authoritative framework for modeling molecular data in a hypothesis-agnostic and discovery-driven way. Second, we assume that a systems approach, linking the longitudinal clinical phenotype to the entire transcriptome, is the least reductionist approach to developing a unified model of inflammation in MOD [24, 54]. And third, we assert that an integrative model provides the best framework for identifying and interpreting novel mechanistic hypotheses regarding pathophysiology, diagnosis, prognosis and treatment of heart failure.

Integrated research strategies have previously been suggested by other groups [28, 55] and successfully implemented by our own [56, 57]. By adopting this strategy, we integrated the current biphasic SIRS/CARS-model into a more unbiased and comprehensive framework. The distinctive immunological features of SIRS/CARS are well-captured by our model at day 1 and 3, when we note the prominent innate immune response and its subsequent suppression and transition to adaptive immunity (Fig. 4B). However, our model has the advantage of incorporating several features that the SIRS/CARS model does not readily accommodate. We note a number of eigengenes that act orthogonally to the innate/adaptive immune response (Fig. 4A), which we functionally classify as metabolic, catabolic, and reparative processes. These processes form a basis for hypotheses beyond the SIRS/CARS framework. For example, it has been previously hypothesized that a key issue to survival is the potential of the patient to initiate recovery strategies, possibly through mitochondrial biogenesis [25]. Recovery mechanisms emerge as distinctive features in our model, and are particularly interesting at later timepoints when the SIRS/CARS framework is no longer helpful (Figs. 4B and 4C). In this way, we functionally characterized distinct stages of recovery (or non-recovery) from MCS surgery, unifying a number of disparate hypotheses. The result is clinically relevant to improving prediction of adverse events and designing early pre-emptive therapeutic interventions.

Finally, while our model was able to accurately identify and integrate many known features of organ dysfunction following MCS surgery, we acknowledge statistical limitations due to the small sample size of our dataset. The small sample size affects the predictive modeling aspect of our study most (particularly the Cox survival models), while the unsupervised approach based on WGCNA has been shown in previous studies to be robust even at small sample sizes (n<30) [58-60]. Furthermore, because of our repeated-measures design, the number of total samples used was sufficiently large when inferring the eigengene network, and when relating it to the clinical parameters using a linear mixed-effect model. To improve reliability of the Cox inferences and the final predictive model, we selected only those genes which where consistently predictive across all timepoints, limiting the influence of statistical fluctuations as much as possible. Thus, most of the statistical limitations in this experiment arise from the heterogeneity of the small patient cohort, rather than the inferential methods used in analysis, which is a limitation that can only be addressed by expanding our scope to a coordinated multi-center study to gain a much larger sample size.

## 5 Conclusions

Our model identifies and synchronizes, at an interpretable level of detail, the many interesting clinical and biological features in the systemic inflammatory response following MCS device implantation. It provides an integrative systems biological model, spanning the full leukocyte genome, while depicting an orchestrated molecular trajectory. This biological sequence of events acts to functionally characterize distinct stages of recovery from MCS surgery, and can help in understanding progressive worsening. This model is also clinically relevant to understanding pre-emptive therapeutic interventions, and to improving prediction of adverse events.

## 6 Competing interests

None of the authors have any competing interests.

## 7 Authors’ contributions

NW helped conceive the study design, conducted the statistical analysis, and wrote the manuscript. GB directed the data acquisition and processing of samples, and helped revise the manuscript. CR participated in the interpretation of results and drafting of the manuscript. JC acquired and processed samples, participated in the interpretation of results and drafting of the manuscript. EC acquired and processed samples, and helped with revision of the manuscript. AE contributed to the drafting of the manuscript and the interpretation of results. MD conceived the study design, participated in the interpretation of results, and drafting of the manuscript. All authors read and approved the final manuscript.

## 8 Additional files

Supplemental files will be linked in the final publication.

## 9 Acknowledgements

We would like to thank Martin Cadeiras for important contributions to the data acquisition, including clinical coordination and consenting of patients. We are grateful to our students Maral Bakir, Ryan Togashi, Giovanny Godoy, Charlotte Starling, Jetrina Maque, Eric Arellano for sample collection and processing. We thank Khris Griffis for double-checking our survival model inferences using nonparametric methods. Finally, we thank our collaborators for useful discussion and comments, including Joe Meltzer, Murray Kwon, Alan Garfinkel, Yina Guo, James Weiss, Elaine Reed, and Peipei Ping.

This research was supported through NHLBI R21 HL 120040-01A1 (Mario Deng), NHLBI R01 HL114437 (James Weiss), NHLBI HHSN26820119999 35C (Peipei Ping), as well as the kind Advanced Heart Failure Research Gifts from Philip Geier, John Tocco, Robert Milo, and Juan Mulder. The funders had no role in study design, data collection and analysis, decision to publish, or preparation of the manuscript.

## 11 Appendix

### 11.1 Organ dysfunction scoring systems

#### 11.1.1 Sequential Organ Failure Assessment (SOFA) Score

The SOFA score is a simple, validated, and widely accepted measure that can be easily obtained using the above parameters, and is frequently used to assess disease severity. It has been shown to be predictive of survival in the critical care unit [14], and has been applied to different study populations. The SOFA score is an integer scoring system that assigns a numerical variable to each of 6 major organ systems to quantify the severity of organ failure. Values range between 0 and 4. Each system’s value is summed into a single SOFA score. Therefore the sum score ranges between 0 and 24, and positively correlates with the severity of the MOD syndrome and clinical outcomes.

Per clinical protocol at our academic center, only those parameters used for clinical assessment were sampled, and only those drugs used for clinical management were included in the database for the study. Therefore, we used the approximate estimation to the SOFA score using the following criteria (Table Supplement 1). The presence or absence of inotropic and vasoactive drugs varies with the SOFA score.

The cardiovascular section of the SOFA score was modified due to certain limitations. Inotropic drugs were presented in the electronic medical records in several different dosage units. Therefore, there was no clear method of comparing the drug doses onto a same scale, and we made the following adjustments:

> 0: Mean arterial pressure greater than 70, No Drugs
>
> 1: Mean arterial pressure less than 70, No Drugs
>
> 2: Mean arterial pressure less than 70, 1 drug, at low dose
>
> 3: Mean arterial pressure less than 70, 1 or more drugs on intermediate doses
>
> 4: Mean arterial pressure less than 70, 1 or more drugs on high doses

The respiratory parameter (PaO2/FiO2 ratio) was frequently missing due to intubation. We therefore made the following replacements in computing SOFA: patients not intubated were given a 1, and patients on respiration support were given a 3.

The neurological parameter (Glasgow Coma Scale (GCS)) was frequently missing from electronic medical charts, due to patients typically being sedated. GCS is a scoring system between 3 and 15 (with 3 being worst, and 15 being best) and is used to determine the conscious status of a patient. It is composed of three parameters: eye response, verbal response, and motor response [61]. Patients are supposed to receive the following SOFA score contributions based on the Glasgow Coma Rating:

> 0: GCS = 15
>
> 1: GCS = 13-14
>
> 2: GCS = 10-12
>
> 3: GCS = 6-9
>
> 4: GCS <6

However, patients who are sedated were assigned GCS scores of 3 (worst), based on the Ramsay Sedation Scale.

#### 11.1.2 Model for End-stage Liver Disease except INR (MELD-XI) score

The MELD-XI score incorporates both bilirubin and creatinine, and is defined [62, 63] by MELD-XI = 5.11 Ln(bilirubin) + 11.76 Ln(creatinine) + 9.44.

### 11.2 Statistical methods

#### 11.2.1 WGCNA

To construct a systems representation, we inferred a weighted co-expression network using the WGCNA package in R [18]. WGCNA efficiently approximates a network adjacency matrix by starting from the cross-correlation matrix. While our dataset has a repeated measures structure, we ignored mixed effects modeling at this step for the sake of computational efficiency, incorporating all samples into the calculation regardless of time or group label. We used the absolute Pearson correlation as an adjacency measure, because we wanted to cluster without regard to the sign of the correlation, in order to improve interpretability under enrichment analyses. WGCNA next alters the adjacency matrix to become approximately scale-free, by raising each element to the smallest exponent that sufficiently maximizes a scale-free fit. The motivation here lies with the assumption that biological networks are approximately scale-free networks [55]. Finally, topological overlap information is used to improve the reliability of the adjacency matrix.

The adjacency matrix was next partitioned by hierarchical clustering, and a dynamic tree-cutting algorithm was used to optimize module assignment. With each module consisting on average of several hundred genes, dimensionality was further reduced using principal component analysis to compute a representative eigengene summarizing the expression of an entire module in a single vector. We computed eigengenes for each module, and used the correlation between eigengenes to define an eigengene network.

#### 11.2.2 Linear Mixed-Effect Model

To relate the modules to the clinical phenotypes, we used a linear mixed-effect model [64]. By using a mixed-effect model, we account for the repeated-measures structure of the data. The statistical significance of the mixed model was corrected for multiple testing using the ‘fdrtools’ R package [65].

#### 11.2.3 Bioinformatics

We computed gene ontology enrichments for each module using the ‘GOsim’ R package [66] (Table Supplement 2). We conducted a pathway analysis on each module using Strand NGS bioinformatics software [20]. Module gene lists were analyzed with a library of “Legacy” pathways from Strand NGS’s Database. We selected the top 5 pathways based on number of entities matched and the p-value. We then used Strand’s natural language processing (NLP) algorithm to identify “pathway hubs,” or the most highly connected genes within each pathway. This algorithm uses IntAct and PubMed abstracts to extract gene interactions, and rank genes based on global connectivity and a relation score. Additionally, we analyzed each module for enrichment of transcription factor binding sites using the whole genome rVista tool [21]. The top genes, pathways, and transcription factor binding sites of interest are indicated in Table Supplement 3.

#### 11.2.4 Cox Proportional Hazards Model

To relate the modules to survival outcomes, we used a Cox mixed-effect model, the ‘coxme’ R package [67]. However, we anticipated significant dynamic effects that could lead to problems with a single model for all timepoints, and also fit separate Cox models at each timepoint using the ‘survival’ R package [68]. When creating the final model, we used the elastic-net implementation in the ‘glmnet’ R package [69].

#### 11.2.5 Principal Component Analysis

To characterize a typical eigengene time course following the surgery, we looked at median eigengene values. Visualization was done using a heatmap (‘heatmap.2’ R package) and PCA biplot (‘biplot’, ‘pca’ R packages). Salient temporal features were found by scaling the medians about their temporal aspect, converting the median eigengene levels to standard z-scores. At each timepoint, module relevance was inferred by then sorting the median eigengenes according to z-score, with the most salient features having the most extreme z-scores.

## 12 Figures

### 12.1 Analysis of clinical parameters

**Figure 1:**
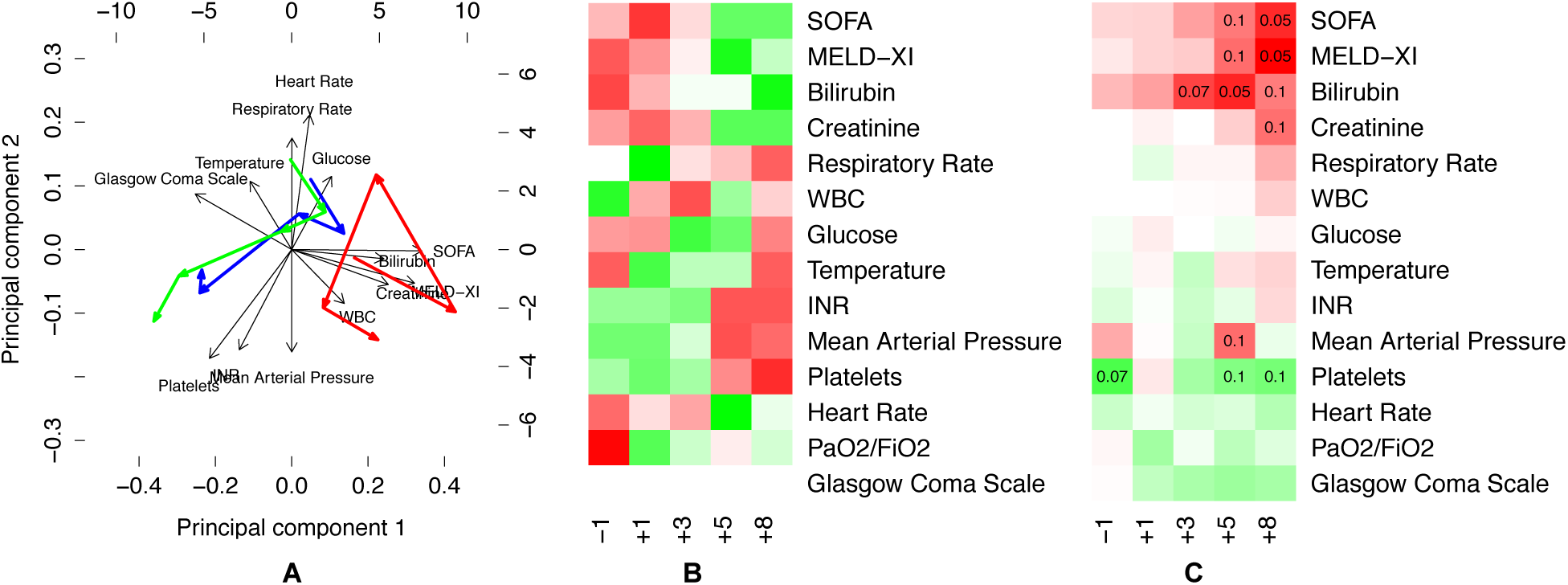
Analysis of clinical parameters. To get a thorough understanding of the phenotype, we analyzed the evolution of clinical parameters over time. **(A)** We projected the timepoint medians onto the leading principal components of the phenotype, and charted the median timecourse (blue) on a biplot. We then separated out the survivors (green) and non-survivors (red). We note a clear separation along the SOFA and MELD-XI dimensions. **(B)** We made a heatmap of the median values for each parameter at each timepoint. Each row is scaled to z-scores to bring out temporal contrasts, where red is upregulated and green is downregulated. On day 1 following surgery, we see the SOFA score peak, the platelet count, temperature, and respiratory rate trough. The white blood cell count reaches a maximum on day=+3, and then recovery occurs on day +5 and +8 as the platelet count rises and the SOFA score decreases. **(c)** We fit univariate Cox models for each clinical parameter at each timepoint, made a heatmap using signed –log p values (where the sign comes from the model coefficient), and displayed corrected q-values below q<0.2. Notice that, with time, the organ dysfunction scores become highly predictive of survival (top right). Platelet count is also moderately predictive of survival, both at later points and before surgery.

### 12.2 WGCNA network

**Figure 2:**
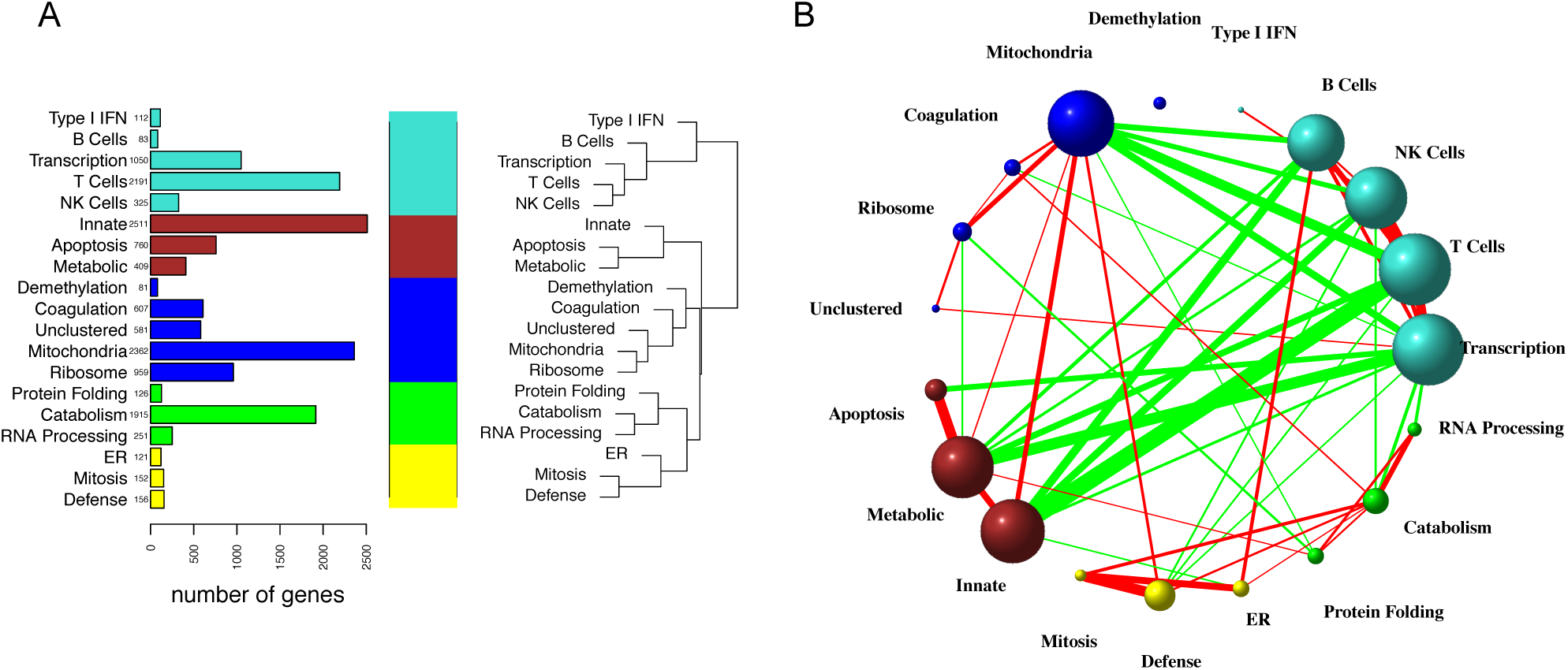
WGCNA network. **(A)** The WGCNA network has 19 modules, and we used gene ontology enrichment analysis to infer module labels. To simplify the representation further, we hierarchically clustered the eigengenes to identify 5 superclusters, and color-coded them with distinct colors. Here, the turquoise supercluster can be understood as the adaptive immune system, the brown supercluster as components of the innate immune system, blue as a metabolic module, green as a catabolic module, and yellow as a reparative supercluster. The 4 most populated modules are T cells, catabolism, innate immunity, and mitochondria, all from separate superclusters. **(B)** WGCNA approximates the eigengene network using the Pearson correlation as an adjacency matrix. We display links after thresholding at r>0.3. The color of each link indicates positive (red) or negative (green) correlation. The width of each link is proportional to the Fisher transformed correlation coefficient. The size of each node is proportional to the eigenvector centrality. Notice the strong negative correlations between the innate and adaptive immune superclusters, the positive correlations of the mitochondria module with the innate module, and the negative correlations of the mitochondria module with the adaptive immune supercluster. The demethylation module is relatively disconnected

### 12.3 Mapping eigengenes to phenotypes

**Figure 3:**
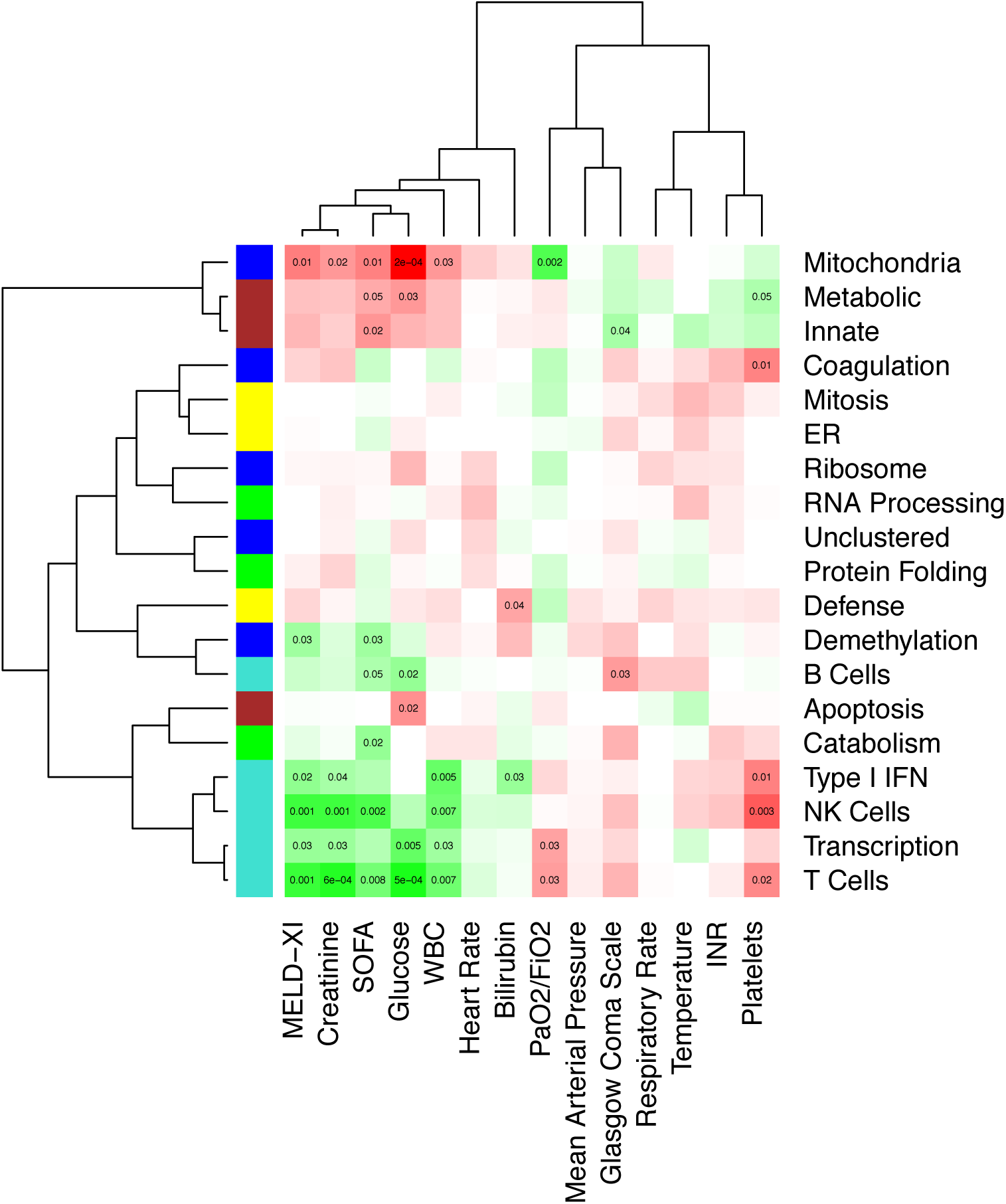
Mapping eigengenes to phenotypes. We inferred relationships between eigengenes and clinical parameters using a linear mixed-effect model, accounting for within-patient variance, and controlling for age, sex, race, diabetes, ischemic etiology, platelet transfusion, device type, plasmapheresis, and immunosuppression. We bi-clustered on the signed -log p-values to identify patterns, while displaying significant q-values. Note that the SOFA and MELD-XI scores are positively correlated with the innate (brown) and metabolic (blue) superclusters, and negatively correlated with the adaptive (turquoise) superclusters. Conversely, platelet count is positively correlated with the adaptive (turquoise) supercluster, while negatively correlated with the innate (brown) supercluster. Finally, the mitochondria module, which relates to both immunity superclusters, is associated with multiple indicators of organ failure.

### 12.4 Time-dependent eigengene analysis

**Figure 4:**
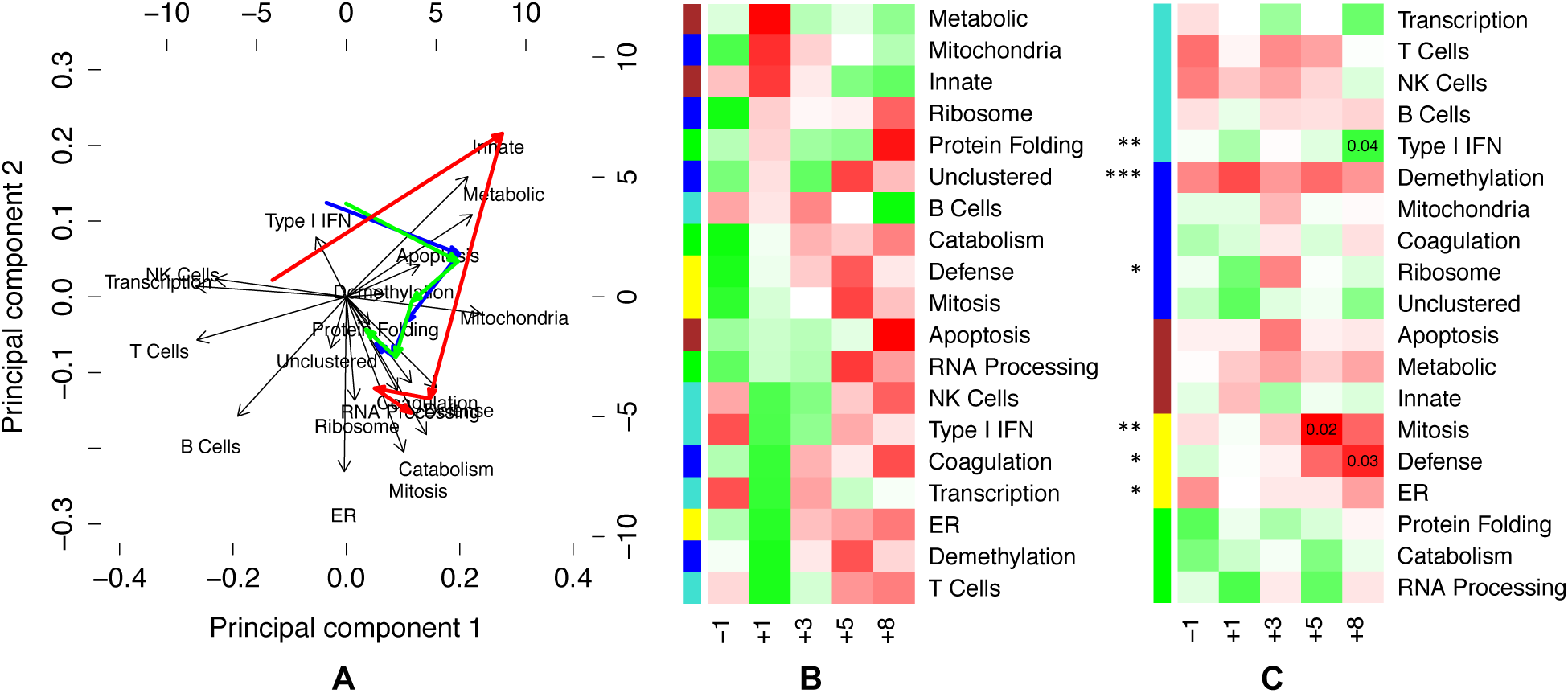
Time-dependent eigengene analysis. To gain a better understanding of the eigengene network, we analyzed the evolution of expression levels over time. **(a)** We projected the timepoint medians onto the leading principal components of the eigengene expression, and charted the median timecourse (blue) on a biplot. We then separated the survivors (green) and non-survivors (red). **(b)** We made a heatmap of the median values for each eigengene at each timepoint. Each row is scaled to z-scores to bring out temporal contrasts, where red is upregulated and green is downregulated. Here, we ordered the rows by sorting z-scores at day 1. **(c)** To infer which eigengenes are related to survival outcomes, we fit a multivariate mixed-effects Cox model to the eigengenes (asterisks), and then fit univariate Cox models at each timepoint (uncorrected p-values displayed), with the heatmap using signed –log p values (where the sign comes from the model coefficient). The mixed-effects Cox model finds demethylation to be highly significant and consistent across timepoints, which we interpret as reflecting that all non-survivors were male (see Discussion). Note also that the reparative (yellow) supercluster emerges as important in the last two timepoints, along with the Type I IFN module.

### 12.5 Module p-value distributions

**Figure 5:**
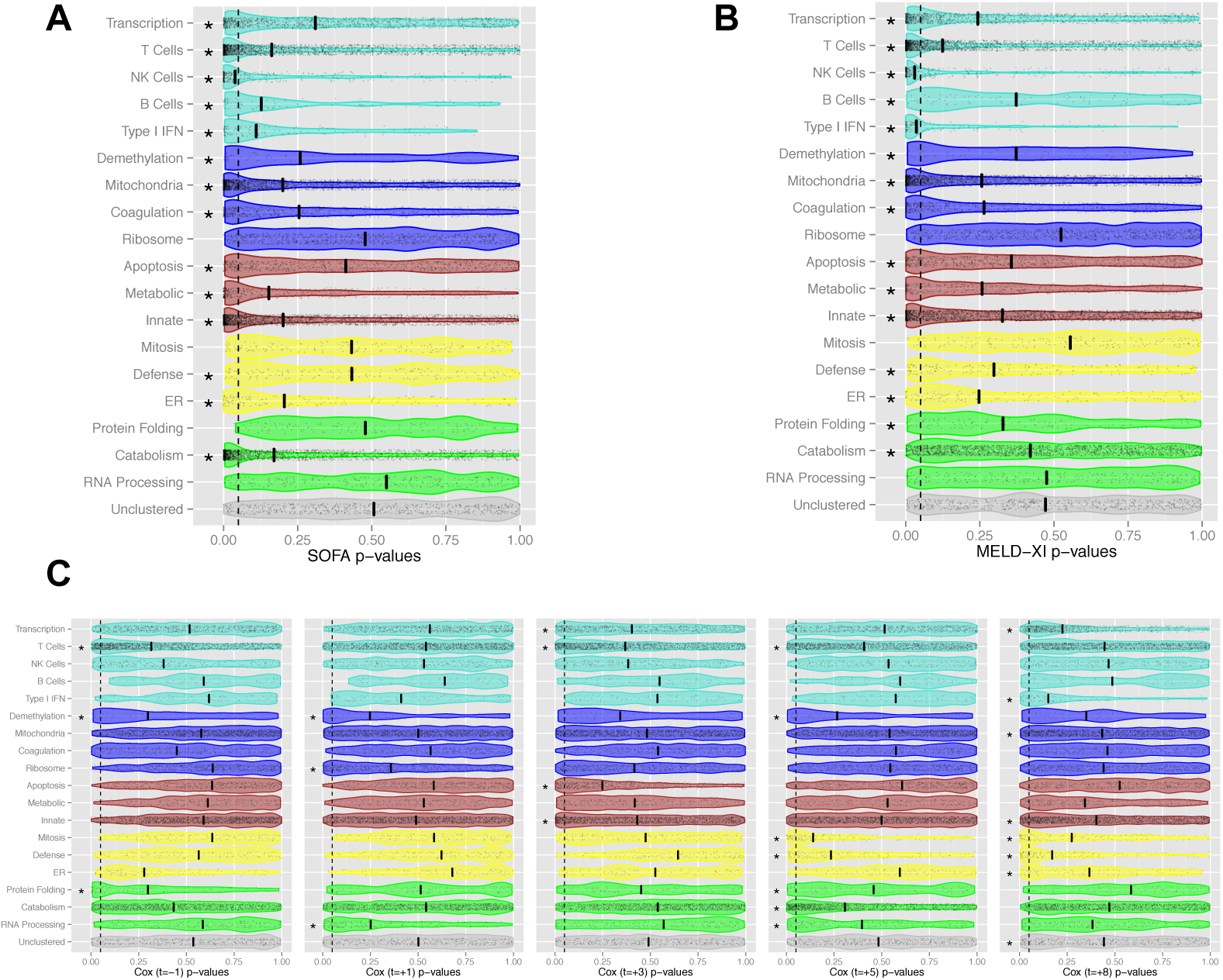
Module p-value distributions. We assessed the p-distributions in each module in relationship to organ failure and survival. We marked the median p-values with a line, and used the binomial test around p=0.05 to assess the balance each p-distribution. Asterisks are shown for statistically significant modules (with Benjamini-Hochberg correction). **(A and B)** We computed p-values for each gene-phenotype association using a linear mixed effect model, and analyzed the p-distributions within each module. Note that almost all modules are enriched with genes associated with organ dysfunction, as measured by **(A)** SOFA and **(B)** MELD-XI scores, illustrating the systemic nature of the syndrome. The innate and adaptive immunity supercluster modules have the most skewed p-distributions. This result is consistent with the linear mixed model analysis of the corresponding eigengenes in Figure 3, which identifies strong correlations between organ dysfunction and the innate and adaptive immune eigengenes. Also note that the B Cell and Catabolism distributions are much more skewed for SOFA score than MELD-XI. **(C)** We computed p-values for each gene-survival association using a Cox model at each timepoint, and analyzed the p-distributions within each module. We note several changes in the p-distributions over time. The demethylation and T cell modules show skew across all timepoints, achieving significance multiple times. Prior to surgery, the protein-folding module has significant skew. Immediately following surgery, the RNA processing and ribosome modules become skewed. On day 3, we see skew in the apoptosis, innate, transcription, and T cell modules. On day 5 and day 8 days, the defense and mitosis modules both become highly skewed. Also on day 5, the catabolism and RNA processing modules become skewed, marking activation of both the reparative (yellow) and catabolic (green) superclusters. On day 8, we see significant skew in the type I IFN and transcription modules, as well as the innate and mitochondria, as the reparative (yellow) supercluster remains activated.

### 12.6 Predictive model

**Figure 6:**
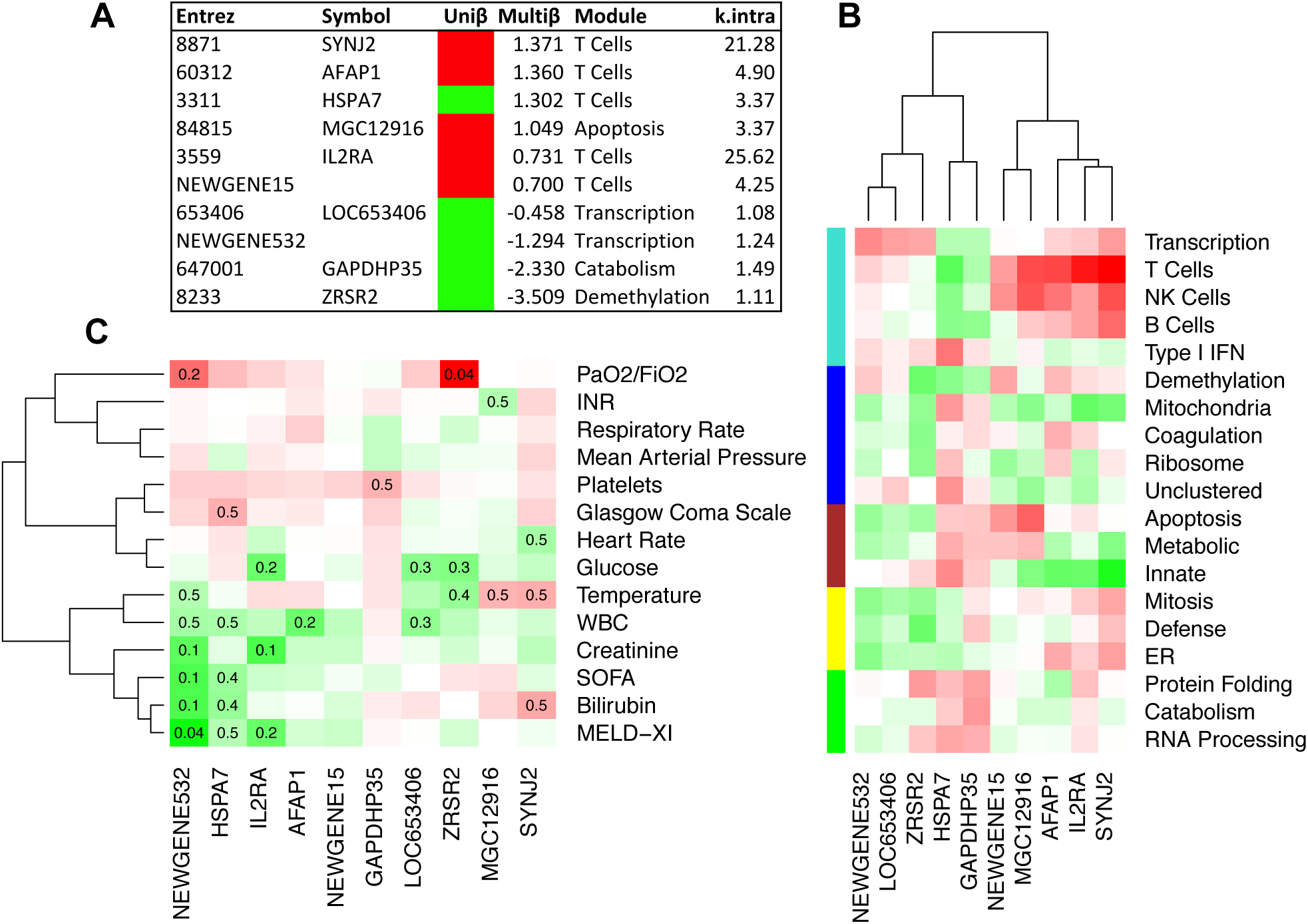
Predictive model. We applied cuts to find genes that had Cox significance level p<0.1 at all 5 timepoints. We then used the elastic net to train a Cox model at timepoint day -1 preoperative. **(A)** We list the genes in the model, and summarize their coefficients and network properties. We distinguished between univariate coefficients (directionality shown in red/green), and the multivariate coefficients (which we report directly), and note that HSPA7 changes directionality under the multivariate model. The Module column lists which module the gene belongs to, and the last column is each gene’s intramodular connectivity. **(B)** We show a clustered heatmap of the module memberships of each gene. Note that SYNJ2, IL2RA, AFAP1, MGC12916 and HSPA7 have strong associations with the adaptive and innate superclusters. **(C)** We bi-clustered the signed –log p-values (the sign comes from the linear mixed-effect model coefficient), and display all Benjamini-Hochberg corrected q-values less that *q*<0.05. Note that NEWGENE532 has very strong associations with the organ failure scores, and ZRSR2 has a very strong association with PaO2/FiO2.

## 13 Tables

### 13.1 Main characteristics of the study samples

**Table 1:** Main characteristics of the study samples.

|  | Characteristic | Total | Survivors | Non-survivors | p-value |
| --- | --- | --- | --- | --- | --- |
| **Gender** | Male | 17 | 12 | 5 | 0.29 |
|  | Female | 5 | 5 | 0 |  |
| **Race** | White or Caucasian | 17 | 13 | 4 | 1 |
|  | Black or African American | 2 | 2 | 0 |  |
|  | Other | 3 | 2 | 1 |  |
| **Age** | Median (IQR) | 62.5 (42-66.5) | 61 (38.5-65) | 65 (62.5-76) | 0.085 |
| **Etiology** | Ischemic | 6 | 2 | 4 | 0.009 |
|  | Non-Ischemic | 16 | 15 | 1 |  |
| **Diabetes** | Diabetic | 7 | 6 | 1 | 1 |
|  | Non-Diabetic | 15 | 11 | 4 |  |
| **INTERMACS** | 1 (Critical cardiogenic shock) | 2 | 1 | 1 | 0.23 |
|  | 2 (Progressive decline) | 12 | 10 | 2 |  |
|  | 3 (Stable, but inotrope dependent) | 7 | 6 | 1 |  |
|  | 4 (Resting symptoms) | 1 | 0 | 1 |  |
| **MCS Device** | HeartMate II | 13 | 11 | 2 | 0.25 |
|  | HeartWare | 1 | 1 | 0 |  |
|  | HeartMate II, CentriMag | 4 | 2 | 2 |  |
|  | TAH | 1 | 1 | 0 |  |
|  | PVAD | 2 | 2 | 0 |  |
|  | CentriMag BVAD | 1 | 0 | 1 |  |

### 13.2 Summary of median timecourse features

**Table 2:** Summary of median timecourse features.

| Time | Upregulated |  | Downregulated |  |
| --- | --- | --- | --- | --- |
|  | Phenotype | Eigengene | Phenotype | Eigengene |
| -1 | MELD-XI, bilirubin, temperature, PaO2/FiO2 | Type I IFN, Transcription | WBC, MAP | Mitosis, catabolism, defense, mitochondria, ribosome |
| +1 | SOFA, creatinine | Metabolic, mitochondria, innate | Respiratory rate, temperature, platelets, MAP, PaO2/FiO2 | T cells, demethylation, ER, transcription, coagulation, Type I IFN, NK cells |
| +3 | WBC | B cells, transcription | Glucose, INR | Type I IFN, NK cells |
| +5 | MAP, INR, platelets | Mitosis, Defense, RNA processing | MELD-XI, SOFA, creatinine, glucose, heart rate | Innate |
| +8 | Platelets, temperature, INR, glucose, respiratory rate | Apoptosis, protein folding, coagulation, catabolism | SOFA, bilirubin, creatinine | B cells, innate, metabolic, mitochondria |

### 13.3 Modified SOFA score

**Table Supplement 1:** Modified SOFA score.

13.3 Modified SOFA score **Table Supplement 1: Modified SOFA score.**
| Score | 0 | 1 | 2 | 3 | 4 |
| --- | --- | --- | --- | --- | --- |
| **Respiration** |  |  |  |  |  |
| Respiratory Status * | -- | No respiratory support | -- | On mechanical ventilator | -- |
| **Coagulation** |  |  |  |  |  |
| Platelets (x1000/uL)<sup>†</sup> | > 150 | < 150 | < 100 | < 50 | < 25 |
| **Liver** |  |  |  |  |  |
| Bilirubin (mg/dL) | < 1.2 | 1.2 - 1.9 | 2.0 - 5.9 | 6.0 - 11.9 | > 12.0 |
| **Cardiovascular** |  |  |  |  |  |
| Mean arterial pressure (mmHg)<sup>‡</sup> | No hypotension | MAP<70 mmHg with no drugs | MAP<70 mmHg, one pressor medication | MAP<70 mmHg, one or more pressor medication at moderate dose | MAP<70 mmHg, one or more pressor medication at high dose |
| **CNS** |  |  |  |  |  |
| Glasgow Coma Scale<sup>§</sup> | 15 | 13 - 14 | 10 - 12 | 6 - 9 | < 6 |
| **Renal** |  |  |  |  |  |
| Creatinine | < 1.2 | 1.2 - 1.9 | 2.0 - 3.4 | 3.5 - 4.9 | > 5.0 |

\* (Modified) PaO<sub>2</sub> and FiO<sub>2</sub> values are not recorded on medical charts for PaO<sub>2</sub>/FiO<sub>2</sub> ratio calculation
† Corrected for platelet transfusion
‡ (Modified) Pressor medication given to patients are presented in several different metric units. Therefore, the CV score was modified into a general categorization of medication dosage level.
§ Neurologic status calculated using the Glasgow Coma Scale (Teasdale *et al*). Sedated patients are given a GCS score of 3 (CNS SOFA score of 4) based on the Ramsay Sedation Scale (Ramsay *et al*).

### 13.4 Gene Onotology Enrichment Analysis

**Table Supplement 2:**
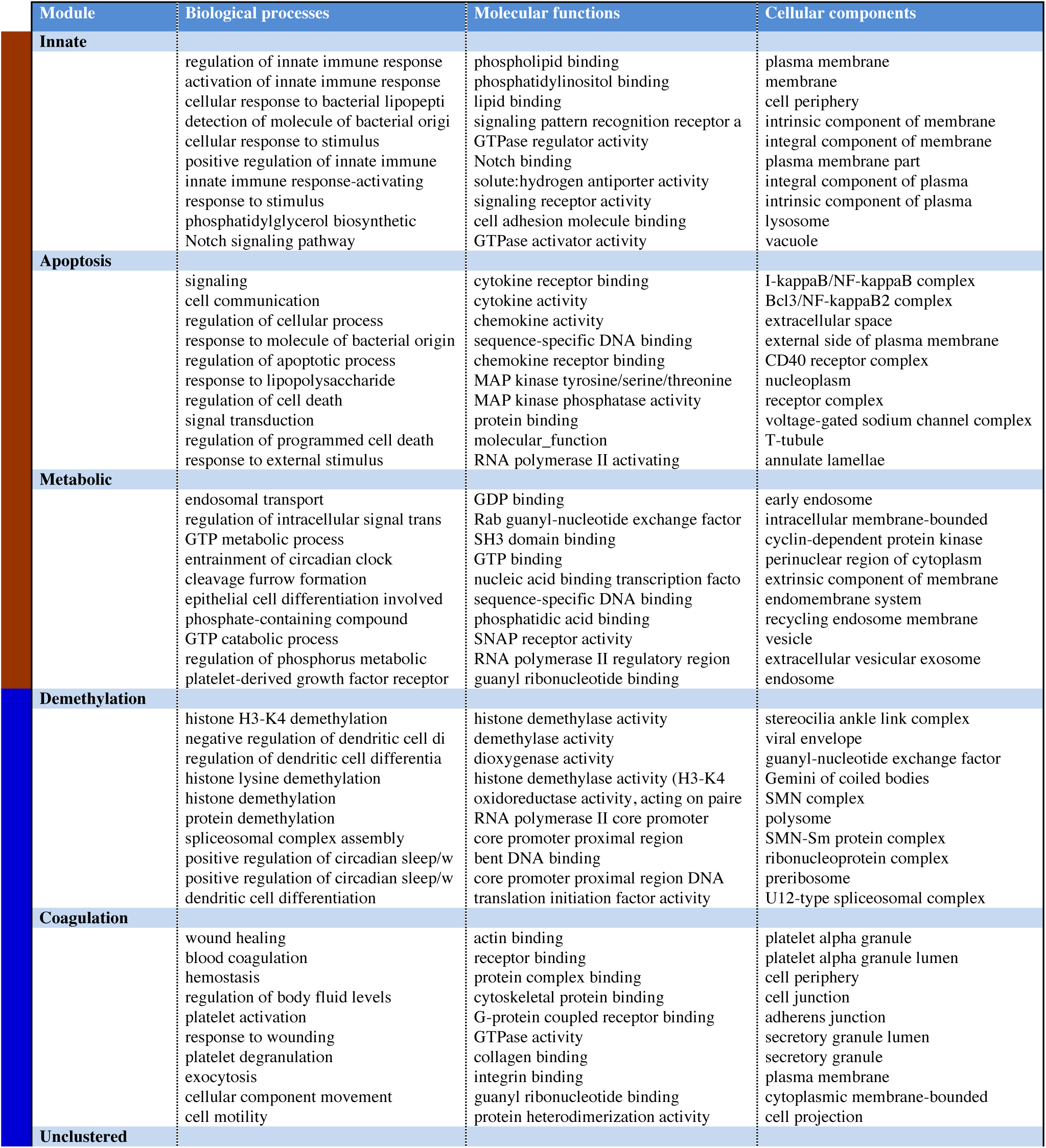
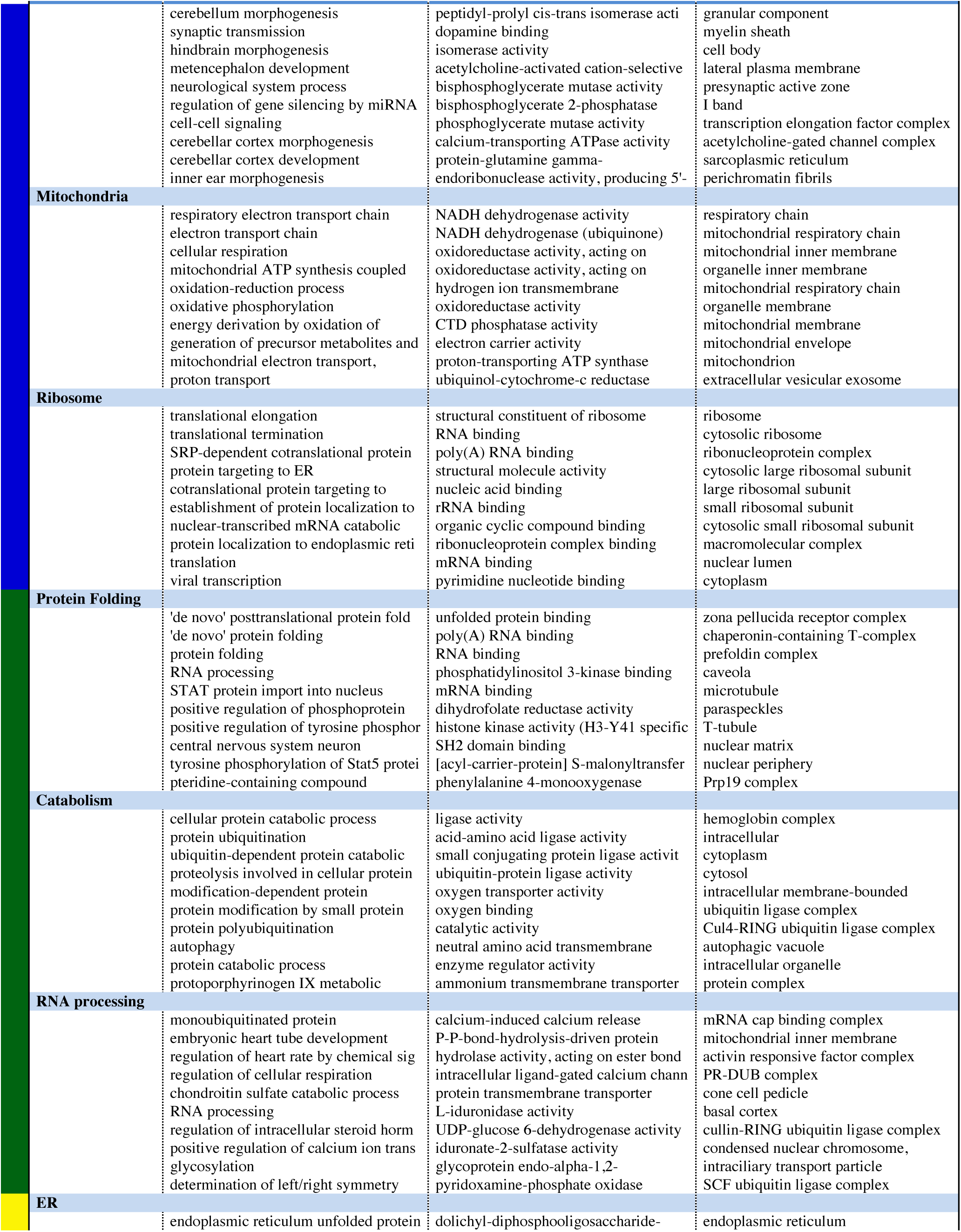
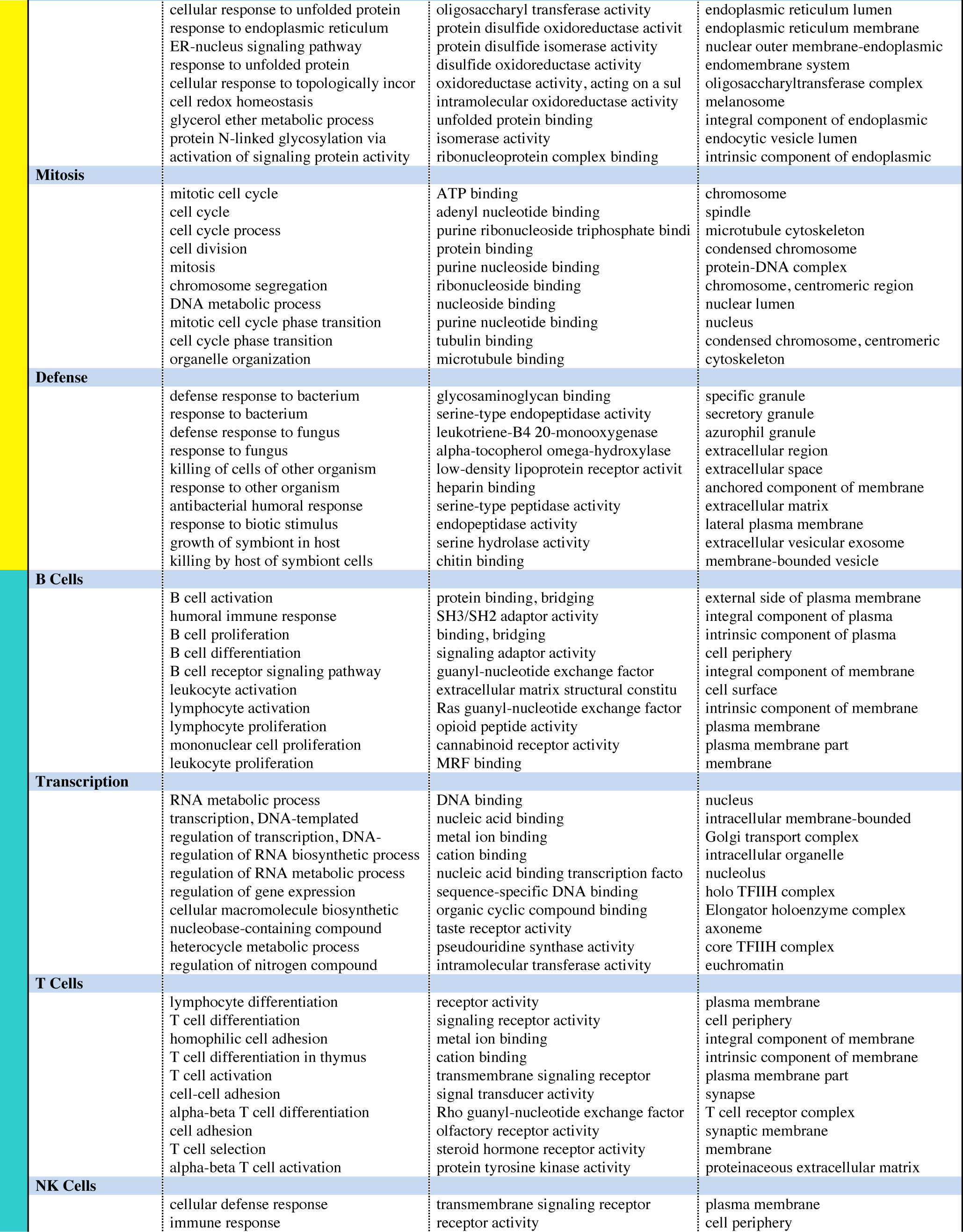
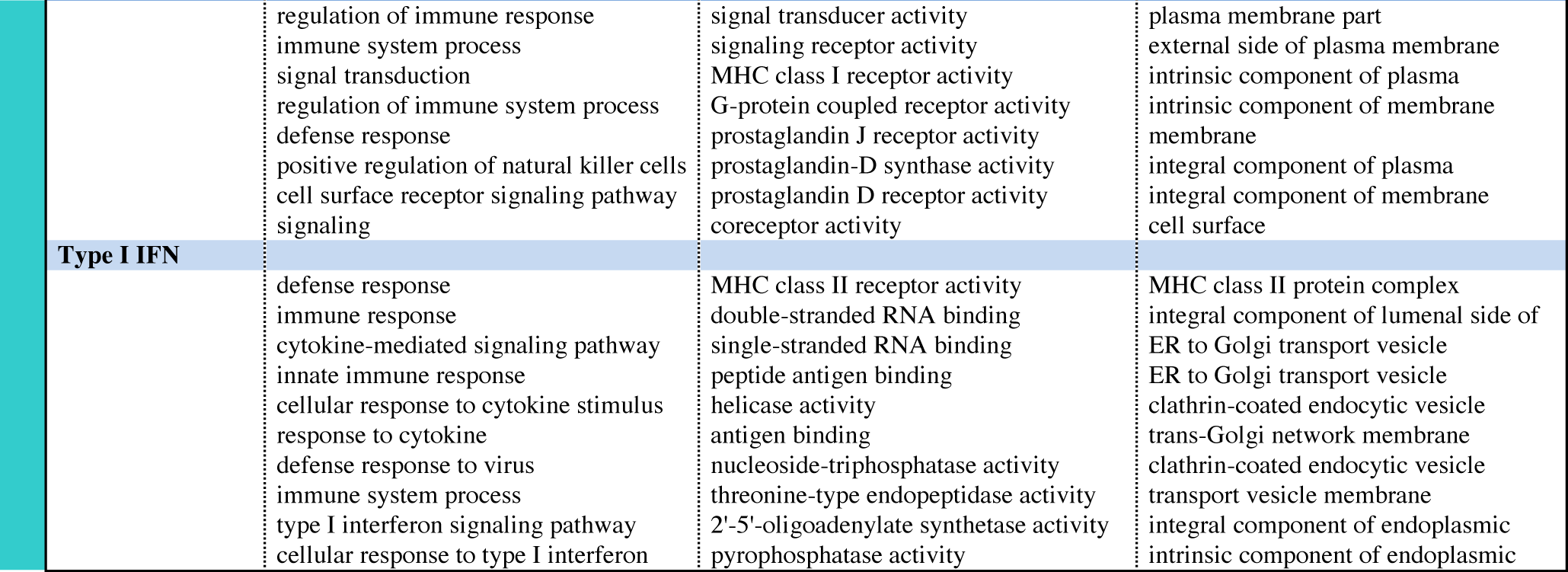
Gene Onotology Enrichment Analysis. We used GoSIM to infer gene ontology enrichments. For each module, we show the top 10 enriched terms, in each of the 3 categories of biological processes, molecular functions, and cellular components.

### 13.5 Pathway analysis and transcription factor binding site enrichments

**Table Supplement 3:**
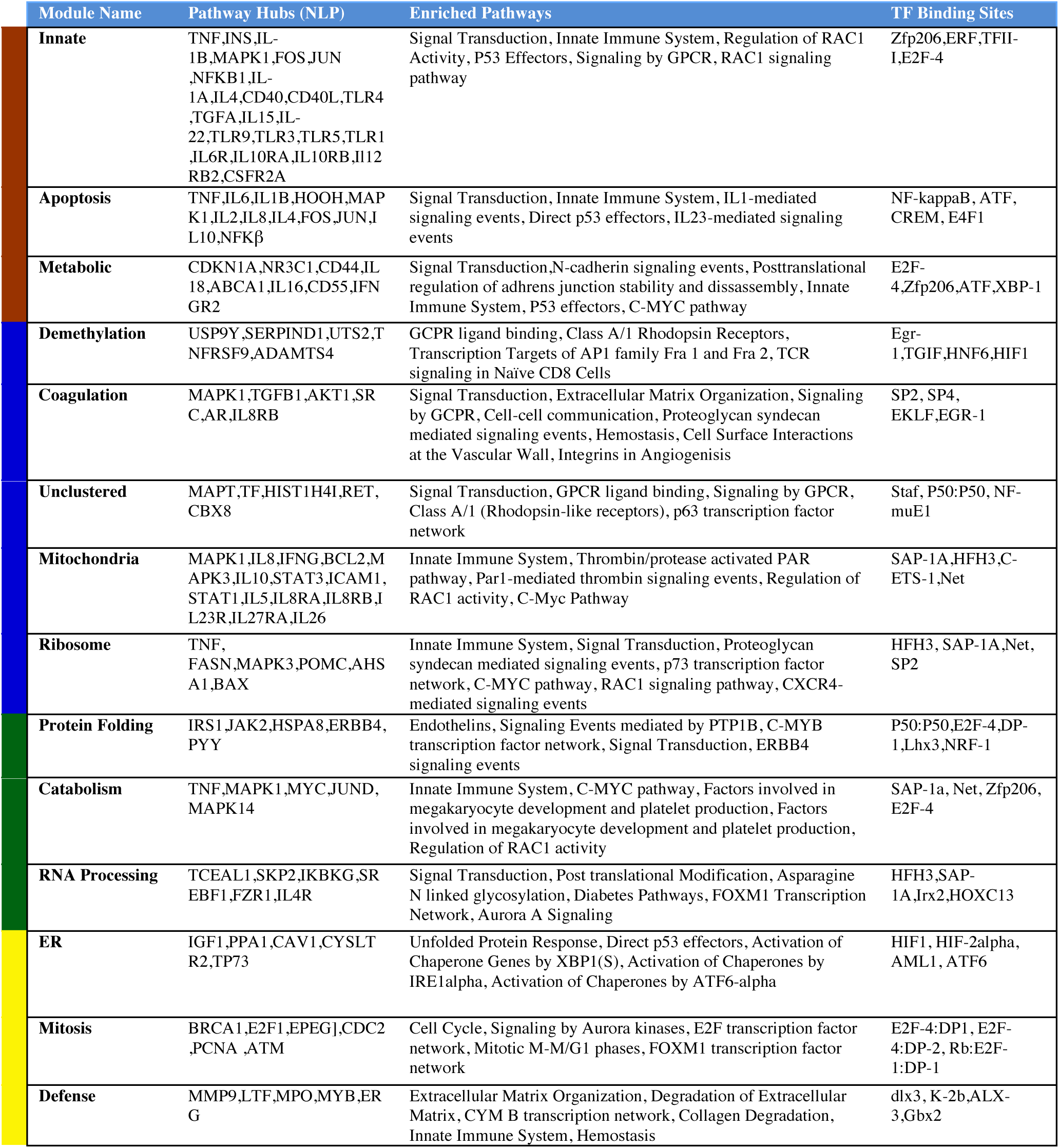
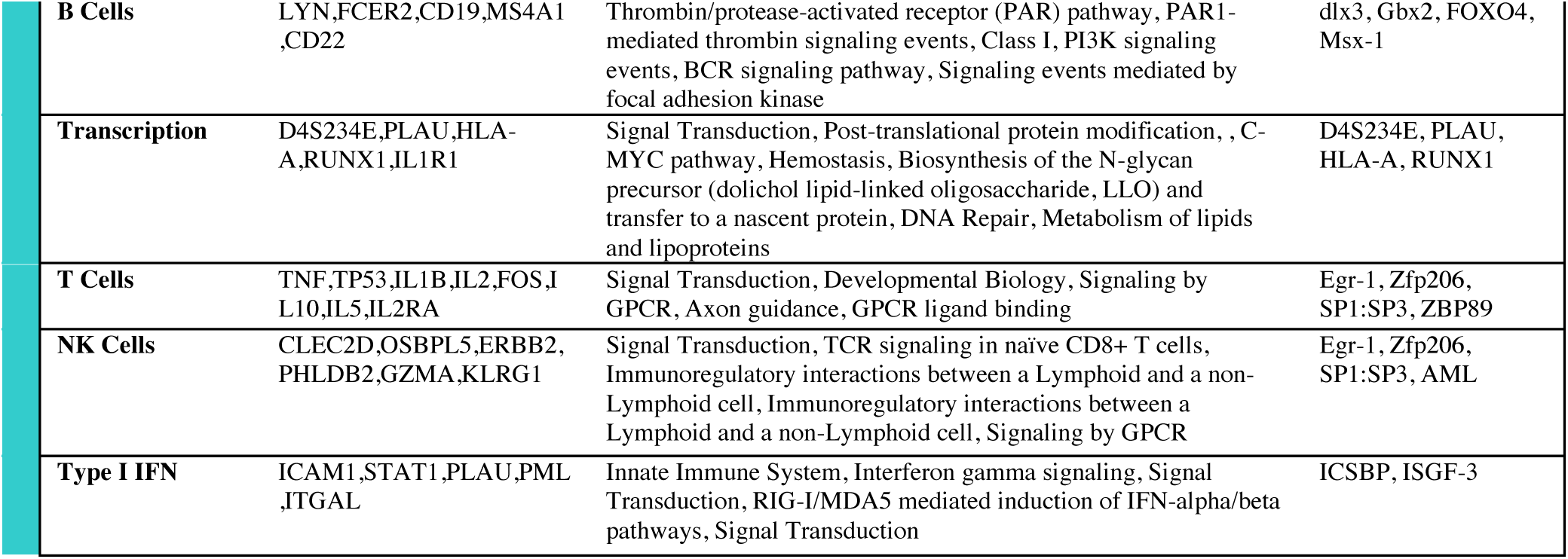
Pathway analysis and transcription factor binding site enrichments. We conducted pathway analyses using Strand NGS bioinformatics software. We identified the top pathways and their hubs for each module. In addition, we used Whole-Genome rVista to analyze enrichment transcription factor binding sites in each module.

